# Avoiding pitfalls: Trace conditioning and rapid aversive learning during route navigation in desert ants

**DOI:** 10.1101/771204

**Authors:** Antoine Wystrach, Cornelia Buehlmann, Sebastian Schwarz, Ken Cheng, Paul Graham

## Abstract

The ability of bees and ants to learn long visually guided routes in complex environments is perhaps one of the most spectacular pieces of evidence for the impressive power of their small brains. While flying bees can visit flowers in an optimised sequence over kilometres, walking ants can precisely recapitulate routes of up to a hundred metres in complex environments. It is clear that route following depends largely on learnt visual information and we have good idea how views can guide individuals along them, however little is known about the mechanisms that control route learning and development. Here we show that ants in natural environments can actively learn a route detour to avoid a pit trap and that this depends on a process analogous to aversive trace conditioning. Views experienced before falling into the trap become associated with the ensuing negative outcome and thus trigger salutary turns on the subsequent trip. This drives the ants to orient away from the goal direction and avoid the trap. If the pit is avoided, the novel views experienced during the detour become positively reinforced and the new route crystallises. We discuss how such an interplay between appetitive and aversive memories might be implemented in insect neural circuitry.

## Introduction

For many different animals, a common navigational mode is to learn habitual, often idiosyncratic, routes to navigate between familiar places of interest (Pigeons: Biro, Meade, & Guilford, 2004; Rats: Calhoun, 1963, Hartley et al., 2003; Ants: Collett et al., 1992; Kohler & Wehner, 2005, Humans: Dee, 2005; Monkeys: Di Fiore & Suarez, 2007; Bees: Woodgate et al., 2017). Routes like these represent an efficient and safe way of navigating through familiar terrain. Probably because they are easy to study in the wild and have reliable navigation behaviour, solitary foraging ants have become a model system for investigations into the sensori-motor and learning basis of visual route guidance as a general animal behaviour (Knaden & Graham, 2016).

Initial exploration in animals can be driven by innate responses to visual or social information (Aron, Pasteels, & Deneubourg, 1989; Collett, 2010; Graham, Fauria, & Collett, 2003; Heusser & Wehner, 2002; von Frisch, 1967). Exploring animals can enter unfamiliar terrain and be safely ‘connected’ to the starting point of their journey by their Path Integration (PI) system (Mittelstaedt & Mittelstaedt, 1980). This method, akin to the basic human sense of direction (Loomis et al., 1993), continuously provides an internal estimate of position relative to the origin of a journey. For ants, PI memories enable individuals to strike relatively straight paths on their early forays between profitable food locations and their nest. It is during these early forays guided by innate strategies that individuals learn the environmental information needed to subsequently guide routes. Indeed, after a few trials, their reliance on learnt environmental information will override their trust in path integration or other innate biases (Collett, Graham, & Durier, 2003; Kohler & Wehner, 2005; Mangan & Webb, 2012; Schwarz, Wystrach, & Cheng, 2017).

We have a good idea how ants can memorise visual scenes, and how these visual scenes can be subsequently used for guidance along a familiar route. Indeed, route following models (Baddeley, et al., 2012, Wystrach et al., 2014; Zeil, 2012; Kodzhabashev & Mangan, 2015) fits well with the ants’ observed behaviour (Collett, Lent, & Graham, 2014; Wystrach et al., 2012; Schwarz, et al., 2017; Wystrach et al., 2014) and recent neuro-computational work sketches how visual route guidance is implemented in the insect brain (Ardin et al., 2016; Webb & Wystrach, 2016). However, little is known about the mechanisms that control route learning and development.

It has been observed that foraging routes are not completely fixed, as they can gradually (Durier, Graham, & Collett, 2004; Wystrach et al., 2011) or abruptly (Mangan & Webb, 2012) evolve. Therefore, studying how ants’ idiosyncratic routes are updated offers an opportunity to understand the underlying learning mechanisms.

Here we used champion route follower solitary foraging ants of two species (*Cataglyphis velox* and *Cataglyphis fortis*) to investigate how naturally encountered consequences (reinforcing or aversive) impacts the development and updating of their routes. Results show that the ongoing refinement of habitual routes depends on an interesting interplay between aversive and appetitive conditioning. Notably, aversive experiences such as getting trapped for a while lead to changes in the valence of the specific visual memories that preceded the negative experience, a process analogous to ‘aversive trace conditioning’ from learning theory (Bouton, 2007). On subsequent runs, these aversive memories trigger salutary avoidance behaviours and expose the ant to new visual scenes which, if the animal is successful in avoiding the trap, are learnt as attractive. Remarkably, this process enables individuals to rapidly reshape routes to avoid large invisible obstacles in their natural environment.

Our approach, which can be qualified as ‘experimental ethology of learning’ (Freas, Fleischmann, & Cheng, 2019; Wystrach et al., 2019), offers an opportunity to understand the interaction between learning mechanisms and real-world complex navigational behaviours.

## Results and discussion

### Ants can reshape their route to circumvent a trap

We let Australian solitarily foraging ants *Melophorus bagoti* shuttle back and forth between their nest and a feeder full of cookie crumbs located 5 m away. For the outbound trip, the ants had to walk through a long and narrow channel suspended 15 cm above the ground that connected the nest directly to the feeder. For the way back to the nest, the ants loaded with a cookie were free to navigate on the desert ground. After a day of shuttling back and forth, all marked ants had established a fairly direct homing route to the nest (Fig 1Ai). We then opened a pit, previously buried inconspicuously into the desert floor, creating a 2 m long, 10 cm wide trap perpendicular to the nest-to-feeder route. During their first homing trial with the trap, all trained ants ran as usual along the first part of the route and suddenly dropped into the trap. The trap was 10 cm deep so that ants could see only the sky. The trap had slippery walls to prevent the ant escaping and contained small twigs, which desert ant naturally tend to avoid as they impede walking. The trap offered a single possible exit formed by a stick leading from the base of the trap to the second part of the homeward route. The time the ants were trapped in the pit varied from one to tens of minutes, but once out, all individuals showed no apparent problem in returning directly to their nest (Fig 1Aii). We let ants shuttle back and forth with the trap open and recorded their paths again after 24h. After such training, several ants (4 out of 14) displayed a new route that circumvented the trap (Figure 1Aiii). We replicated these conditions at a larger scale (8 m route and 4 m wide trap) with North African desert ants from Tunisia (*Cataglyphis fortis)* and obtained similar results (Fig 1Bi,ii,iii).

**Figure 1.**
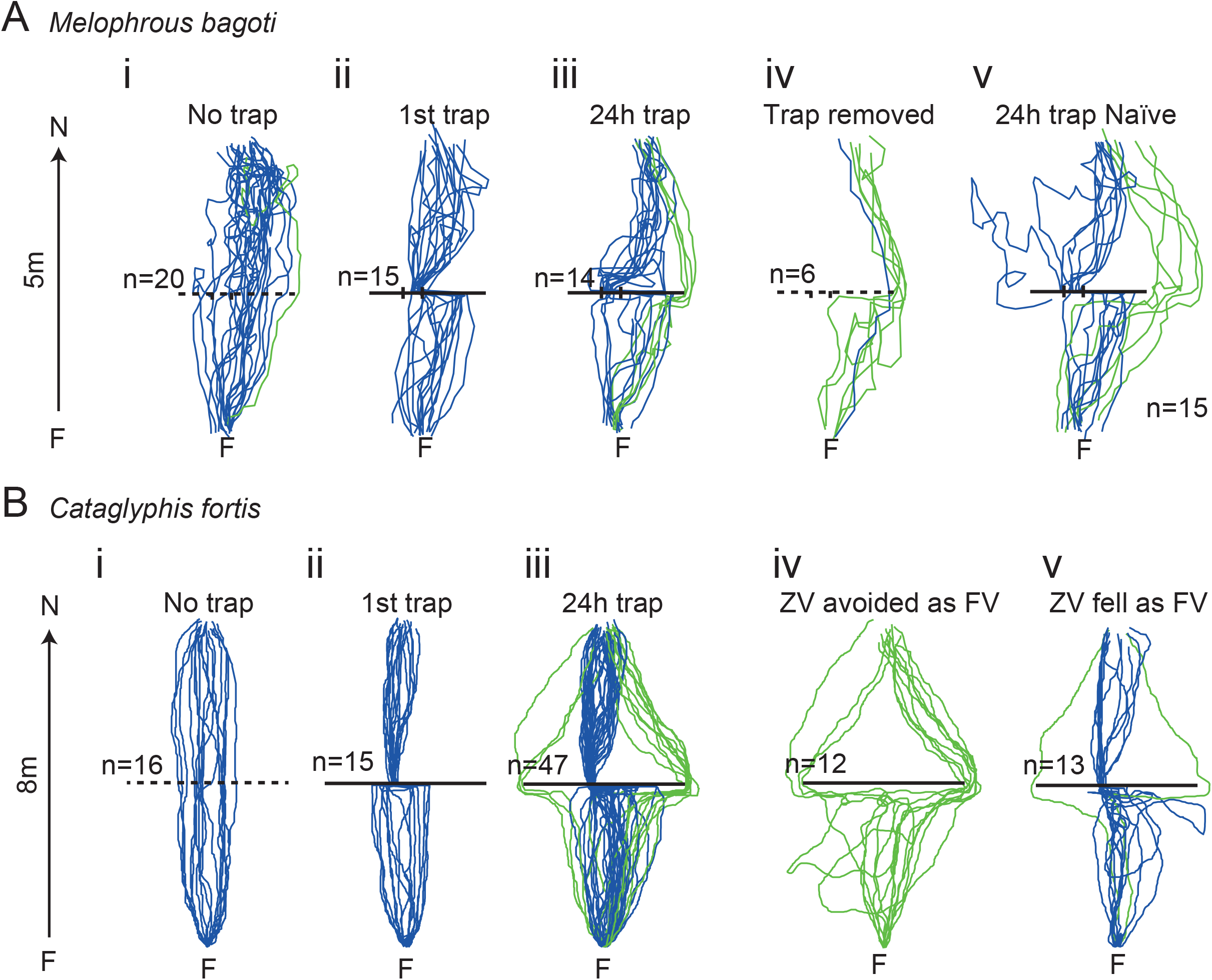
Negative experience shapes ants’ routes. **A.** Australian desert ants (*Melophorus bagoti*) or **B** North African desert ants (*Cataglyphis fortis*) were followed for a series of homeward routes from a permanent feeder (F), with a pit-fall trap in place (solid line) or covered over (dashed line). (i) Control routes of ants between feeder and nest. (ii) The first route after the installation of the pit trap. (iii) Paths after the pit trap has been in place for 24 hours. Green paths from ants that circumvent the pit trap and blue paths from ants that fall into the trap. (iv) and (v) show different conditions for the two species. (Aiv) paths of *M. bagoti* ants that had learnt to circumvent the trap when it was subsequently covered again. (Av) the paths of ants that had begun their foraging life with the trap in place and had 24 hours of foraging experience. (Biv, v) paths of *C. fortis* ants that had learnt to negotiate the trap when tested as zero-vector ants and paths of ants that had avoided (green) or fallen (blue) into the trap, respectively, as full-vector ants.

Why some ants did not learn to circumvent the trap may be due to different reasons. A good proportion of those ants did show modification of their routes by learning to avoid the trap using alternative strategies such as jumping directly onto the exit stick (see red paths in SI1), or simply learning a quick route through the trap by systematically falling in at the same spot and quickly reaching the exit stick with very little search. Finally, some ants did simply not learn, perhaps because they performed too few training trials within the 24h period. In any case, a simple categorisation of whether the ants circumvented the trap or not is sufficient to show that such an effect is unlikely to happen by chance (First_trial_with_pit vs. 24h_with_pit: N_(circumvented)_ /N_(all ants)_ : 0/30 vs. 17/61, Fisher’s Exact test p<0.001), and that there was no apparent difference in detour success between species (24h_with_pit *M. bagoti vs. C. fortis*: N_(circumvented)_ /N_(all ants)_ : 4/14 vs. 13/47, Fisher’s Exact test p=1).

### New routes are based on learnt terrestrial cues

Desert ants are well known to follow habitual routes guided by learnt terrestrial cues although they also have access to their Path Integration (PI) system at all time (Knaden & Graham, 2016), a navigational strategy that is particularly pronounced in *C. fortis*. We carried out several manipulations to demonstrate that learnt terrestrial cues, rather than PI, were controlling the new routes of our foragers.

Ants captured just before entering their nest and then re-released at the feeder are called zero vector (ZV) ants because their PI state is zero, and thus no longer provides correct homeward information. Such ZV ants that had circumvented the trap during their previous (full vector, FV) run were equally successful in their subsequent ZV run (Fig 1Bv) whereas ants that had fallen into the trap as FV ants still did so as ZV ants (previous_FVcircumvented vs. previous_FVfell: N_(circumvented)_ /N_(all ants)_ : 12/12 vs. 2/13, Fisher exact test p<0.001), showing that guidance along the newly learnt route does not requires PI. Note that 2 out of 13 ants fell as FV but avoided the trap as ZV, which further supports the hypothesis of visual route learningWe know that the directional dictates of PI and learnt visual guidance are integrated, even when pointing in different directions (Collett, 2012; Legge et al., 2015; Wystrach, Mangan, & Webb, 2015). Thus, in FV ants, the PI vector points directly to the nest and thus may bias the path towards the trap. Therefore, the paths of ZV ants are more representative of the route as guided by terrestrial cues.

We further tested whether ants that had learnt a new route to circumvent the trap would still follow it, if the trap was removed again. Five out of the six individuals tested displayed the usual detour even though the trap had been removed (Fig 1Aiv; No_pit_initially vs. pit_removed_again: N_(circumvented)_ /N_(all ants)_ : 1/20 vs. 5/6, Fisher exact test p<0.001). This confirms that the detour does not depend on perceiving the trap.

We can further rule out the use of chemical trails, scent marks or other social information, although their use is unlikely in these highly visual solitary foraging ants, by simply observing the idiosyncrasies of the ants’ individual routes, typical in these species (Collett et al., 1992; Kohler & Wehner, 2005; Mangan & Webb, 2012). Routes from an individual ant are remarkably similar across subsequent trials (Fig 2A, Fig S1) but they vary substantially across individuals (Fig 1, SI), showing clearly that ants were using private information.

**Figure 2.**
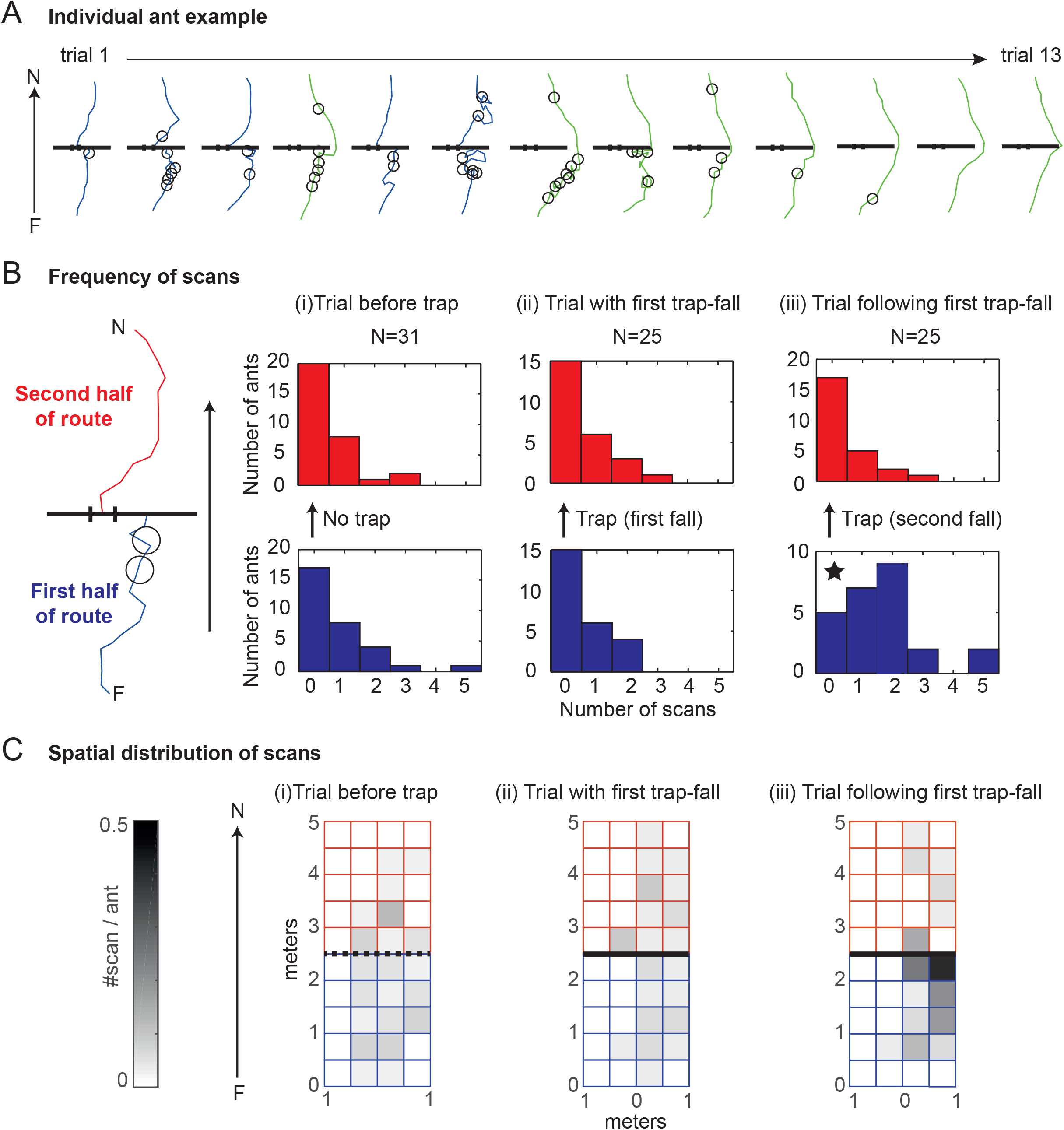
Negative experience modifies the memory of the views experienced before the trap. Individually marked ants were tracked for a sequence of runs before and after the activation of the pit trap. **A.** Sample sequence from a single ant. Locations where the ant stops and scans the world are marked with a circle. As elsewhere in the paper, paths are coloured coded: blue for ant paths that fall into the trap and green paths that avoid the trap. **B.** For each ant the number of scans was recorded before (bottom, blue) and after (top, red) the pit across three focal trials. (i) Trial before the trap was activated; (ii) Trial with first pit-fall; (iii) Trial following first trap-fall. From this we calculated the probability of an ant scanning, and the mean number of scans, for each trial and route segment. Before the pit: (i) N=31 proba=0.35 Mean=0.52±0.85 (ii) N=25 proba=0.40 Mean=0.60±0.87 (iii) N=25 proba=0.40 Mean=0.60±0.87. After the pit: (i) N=31 proba=0.40 Mean=0.60±0.87 (ii) N=25 proba=0.40 Mean=0.56±0.77 (iii) N=25 proba=0.80 Mean=1.64±1.35. **C.** For the same three focal trials, the location of scans is shown relative to the Feeder (F, (0,0)), Nest (N, (0,5)) and Pit Trap (Black line, y=2.5). Darker areas represent higher scan numbers.

Finally, we recorded a cohort of ants that had started their foraging life while the trap was already in place. We did not control how many trials each ant produced but within a period of 24h we observed that several individuals learnt routes that circumvented the trap (Fig 1Av).

The proportion of ants that circumvented the trap was similar between these ants and ants that had some route experience before the trap was set in place (24h_with_trap_naive vs. 24h_with_trap: N_(circumvented)_ /N_(all ants)_ : 5/15 vs. 4/14, Fisher exact test p=1). This shows that a route that circumvents a hidden trap will develop naturally, whether the trap is novel or has always been there.

Taken together, it seems clear that the novel routes displayed by ants were guided by learnt terrestrial cues. The nature of this learnt information is not crucial for our purpose here, but based on past evidence with these species, we can be confident that it is mostly visual (Wehner, 2009). To ease the reading, we will now refer to these learnt terrestrial cues as ‘familiar views’.

### How do ants reshape their routes? Evidence for aversive trace learning

To shed light on the processes that lead from an established route to a new route that circumvents the trap, we tracked all successive trials of individually marked *M. bagoti* ants from the first time they encountered the trap onwards. In addition to their paths, we recorded the locations where ants stopped and scanned their surroundings. Scanning is a typical and obvious behaviour in this species: the navigating ant suddenly stops and rotates on the spot, pausing in different directions successively (Wystrach et al., 2014). Scans are triggered when the ant experiences a decrease in visual familiarity, when multiple directional cues are put into conflict, when running a route has not resulted in success, or simply when naïve ants exit their nest for the first time (Fleischmann et al., 2017; Jayatilaka et al.,2018; Wystrach et al., 2019; Muller & Wehner, 2010). In other words, the occurrence of scanning reflects a directional uncertainty in an ant’s navigational system (Wystrach et al., 2014).

As described above, on the first run with the trap in place, ants rush along their direct homeward route and fall into the trap. Routes of most ants, however, then change from trial to trial and some ants eventually circumvent the trap. It could be that the negative event of falling into the trap triggers learning oriented behavioural routines that occur immediately after the negative event. When ants emerged from the trap, however, they rushed towards their nest as usual, displaying neither more scanning or meandering than before the trap was set (Second part of the route: *Trial before trap* vs. *Trial with first trap fall*: mode = 0 scan/ant in both cases, glme #Scan: z=0.398 p=0.916; glme meander z=0.067 p=0.997; Fig 2Ai and 2Aii, see also Fig 1Aii and Bii)).

An alternative strategy would be to keep traces of the scenes experienced immediately before falling into the trap, and change their valence given the current negative experience of being trapped. In our paradigm, this would predict that ant behaviour will be affected on the run subsequent to falling in the trap when in the area immediately before the trap. Indeed, this is what we observed. Ants displayed a clear increase in scanning behaviours in the region before the trap (mode = 2 scans/ant, Fig 2Biii), significantly more than they had on their previous run (mode = 0 scan/ants, Fig 2Bii) before falling into the trap for the first time (First part of the route: *Trial with first trap fall vs. Trial following first trap-fall*: glme #Scan: z=3.471 p=0.001). Similarly, their paths showed significantly more meandering as they approached the trap for the second time compared to their previous run (First part of the route: *Trial with first trap fall vs. Trial following first trap-fall*: glme meander: z=3.180 p=0.004).

In line with current models of insect associative learning, the experience of the trap triggered a change in the valence associated with the familiar views experienced just before falling into the trap and this classical conditioning of navigational memories leads to distinct behavioural changes on subsequent trips. We consider the observed behaviours to stem from trace conditioning because the views that triggered adjustments of behaviours were those experienced some time before falling into the trap (see Figs. 1 and 2). Mechanistically, this means that the ant must always maintain a temporary trace of the CS to associate with subsequent US, the most important characteristic of trace conditioning.

### Mechanistic implications

The current understanding of visual route navigation is that when familiar views are perceived ants inhibit turning and stimulate forward movement (Ardin et al., 2016; Baddeley et al., 2012), while unfamiliar views would trigger turns (Kodzhabashev & Mangan, 2015). In this model familiarity is synonymous with a positive valence. Our results suggest, however, that being trapped can trigger a punishment signal, so that familiar views become associated with a negative valence. Negative valences trigger turning and scanning behaviours, which will bias the navigator towards other directions. If the new route leads to no further aversive events, and the ant arrives home in good time, the novel views experienced will be positively reinforced and a new visually guided route detour will crystallise. What constitutes the positive reinforcement during route learning is still unclear. It could be reaching the nest or running down the PI accumulated home vector (Ardin et al., 2016). In both cases, it is likely that avoidance behaviour triggered by trace conditioning is then supported by appetitive learning based on positive reinforcement (Fig 3).

**Figure 3.**
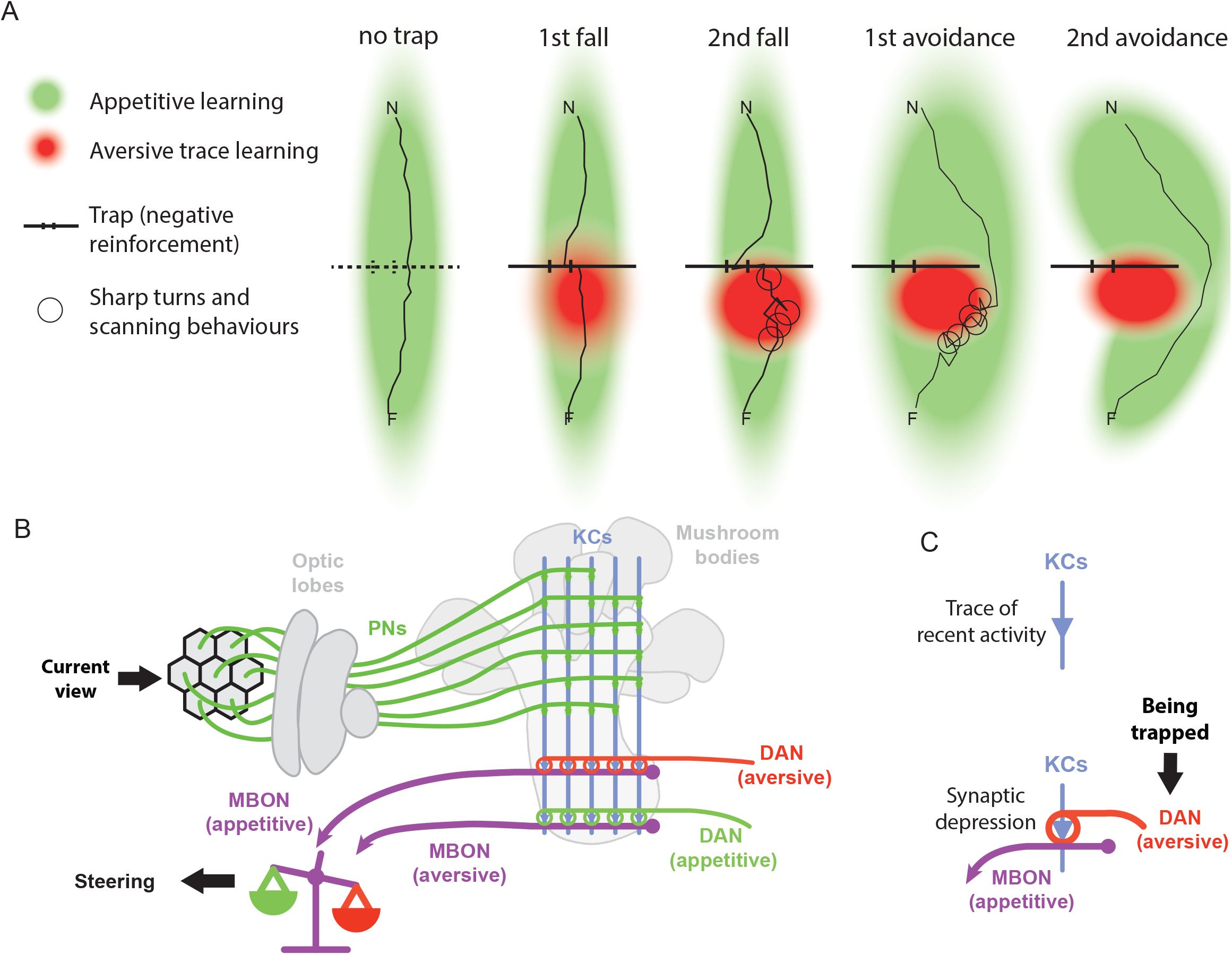
Overview of the appetitive and aversive learning experiences. **A.** Across a sequence of journeys for a typical ant we show the regions of operation for aversive (red) and appetitive (green) conditioning. The aversive region is first formed by trace conditioning on the trials where the ant encounters the trap. Over time a detour develops, and the new route is maintained by appetitive processes. **B**. Schematic of a Mushroom Bodies neural architecture derived from the insect literature. Visual information is sparsely projected via Projection Neurons (PNs) to the Kenyon Cells (KCs). KC activity thus represents views that can be associated with the Mushroom Body output neurons (MBON) mediating appetitive or aversive valences. Such associations result from the modulation of KC-to-MBON synapses, the modulation is generated by the co-activation of KCs and dopaminergic neurons (DAN) relaying the valence of the current situation. The resulting balance between aversive and appetitive MBON activities can then control steering. **C**. The current study suggests trace conditioning as a mechanism to explain the reshaping of routes. First, a trace of the KC activity must be kept for a few seconds (top panel). Second, the co-activation of a dopaminergic neuron modulates the KC-to-MBON synapses of these recently activated KCs (bottom panel). Note that modulation consists of a synaptic depression, which explains why DANs of a given valence modulate MBONs of the opposite valence. Thus an aversive situation, such as being trapped, will be mediated by a DAN decreasing the connection strength between the recently activated KCs and the appetitive MBON. These KCs will no longer activate the appetitive MBON, but still activate the aversive MBON. In other words, the view experienced before the trap will henceforth trigger an aversive response.

Trace learning implies certain neural requirements: 1) Some neural activity must be specific to the views experienced along the route; 2) A trace (or tag) must remain in such view-specific neurons (or their output synapses) after they have fired; 3) A negative event must be able to change the valence associated with the tagged neurons long-term or at least as long as it takes the ant to perform another trial; 4) This change of valence must change the ant’s behaviour when the views are experienced again, in this case triggering turns and scans.

We now have a good idea of the neural underpinnings learning in insects from studies of the Mushroom Bodies (MB), which are assumed to be the seat of the route visual memories (Fig 3; Webb & Wystrach, 2016). The knowledge of the MB circuits gives us direct mapping to the above requirements: 1) Each view experienced can be represented by a specific pattern of activation of Kenyon Cells (KC) in the MB (Ardin et al., 2016); 2) Each KC projects onto multiple output neurons (MBONs) conveying different valences (Aso et al., 2014; Aso & Rubin, 2016). MBONs can therefore be used to tag the outputs of KCs at the synaptic compartments between KCs and specific MBONs. As yet it is unclear where exactly the tagging may happen, as several types of neurons project to these compartments (Liu & Davis, 2009; Perisse & Waddell, 2011); 3) Each KCs–MBON compartment receives modulatory input from specific dopaminergic (or octopaminergic) neurons. These neurons typically convey reward or punishment signals that modulate the KCs–MBON synapses (Cohn, Morantte, & Ruta, 2015), and can lead to long-term memory (Bouzaiane et al., 2015; Isabel, Pascual, & Preat, 2004); 4) KCs–MBON synaptic changes typically modulate an attraction versus avoidance balance for learnt stimuli (Bouzaiane et al., 2015; Cohn et al., 2015).

Route learning seems therefore to fit this picture of MB function remarkably well. By using both appetitive and aversive MBONs for experienced views which activate unique patterns of KCs, views can be tagged and subsequently associated to either valence depending on the type of reinforcing signal. Known neurobiological underpinnings of associative learning in insects can support our behavioural account of aversive trace conditioning in navigating ants.

## Conclusion

We have demonstrated here an ecologically relevant example of trace conditioning, where a negative experience labels specific locations in space that precede an aversive event. Behaviourally, this allows an ant to solve a navigational problem via adaptive route reshaping behaviours on subsequent journeys through the world. More generally, this suggests an interesting interplay between aversive and appetitive visual memories, and between avoidance learning (a form of negative reinforcement) and positive reinforcement, which matches well our current understanding of insect learning circuits.

## Materials and Methods

### Species and field sites

Two species were tested in this study: the Australian red honey ant, *Melophorus bagoti* and the Saharan desert ant *Cataglyphis fortis.* Both species are highly thermophilic ants (Wehner, 1987; Christian & Morton, 1992) that forages solitarily on dead insects and plant materials (Muser et al., 2005). The field site for experiments with *M. bagoti* was located ~10 km south of Alice Springs, Australia, on the grounds of the Centre for Appropriate Technology, in a semi-arid desert habitat characterised by red soil, grass tussocks, bushes, and trees of Acacia and Eucalyptus species. Field experiments with the *C. fortis* were performed in a flat salt pan (34.954897 N, 10.410396 E) near the village of Menzel Chaker, Tunisia.

### Experimental set-ups

The experimental set-up in both species were similar, with a larger scale for *C. fortis* to reflect their typically longer range of foraging. Measurements below are given for *M. bagoti*, followed by *C. fortis* in brackets. Experiments were undertaken with a nest located in an area cleared of grass but surrounded by bushes and trees (or artificially added large black cylinders for *C. fortis*) providing rich visual information for navigation.

In both experimental set-ups, ants moved between their nest and a feeder full of cookie crumbs 5 m (8 m for C*. fortis*) away. The ants' nest was covered with an overturned bucket that had the bottom removed, whose opening at ground level was connected to a straight outbound channel. Ants on their outbound trip were thus channelled from the nest into a 5m long, 10 cm wide and 10 cm high (8 m long, 7×7 cm for *C. fortis*) channel elevated 15 cm above the ground that suddenly ended above the feeder, into which ants would drop. The feeder was a small plastic container sunk into the ground providing biscuit crumbs *ad libitum*. To return to the nest, the ants climbed out of the feeder using a small ramp and walked on the desert ground back to the nest. Removing the feeder ramp at critical times allowed us to control which ants ran their homeward journey and when. Halfway along their homing route, a plastic channel was buried inconspicuously into the desert floor, creating a 2 m long, 10 cm wide (4 m long 7 cm wide for *C. fortis*) trap perpendicular to the nest-to-feeder route. The trap was buried entirely so as to remain invisible to the ants. The trap had smooth walls and was filled with twigs to hinder ant movement. Ants could leave the trap by using a single exit formed of a stick bridge 2 cm wide, connecting the bottom of the trap to the second part of the route. A grid of lines (mesh width: 1 m) was set up by winding strings around pegs (or by painting on the ground with *C. fortis*) and the ants’ homing paths before and after introducing the trap were recorded on squared paper. During initial training the pit was covered by a thin board, with desert sand scattered on top, so that the ants could shuttle unimpeded.

### Experimental protocols

To ensure that ants were experienced before the trap was set, ants that arrived at the feeder were marked with a dot of day-specific enamel paint. Only ants with at least 24 hours experience were recorded. Once the trap was set, the ants’ first homing paths after trap introduction were recorded as well as their paths 24 hours later.

With *M. bagoti*, an additional treatment was enacted. Successful ants that circumvented the trap were marked and, once they return to the feeder again, tested with the trap covered again (as in the initial training).

With *C. fortis* a group of ants were recorded twice. Here, the ants performed their homing route and just before they entered the nest they were taken and released again at the feeder as zero-vector ants.

Another condition was tested with naïve *M. bagoti* ants. The trap was set in place and all ants were marked for 5 consecutive days. After this period, all unpainted ants reaching the feeder were considered ‘naïve’ and were painted with a specific colour. Naïve ants were free to forage for 24h before being recorded.

Finally, a third batch of *M. bagoti* ants were marked with individual colour codes in order to obtain a record of the evolution of individual routes. In this treatment, we recorded both the path and the occurrence of the scanning behaviours typically observed in this species.

### Analyses

Paths were digitised using the software Graphclick. Meander was calculated as the mean angular deviation in direction between successive 30 cm chunks of the ants’ paths. For the ‘*Avoid vs. fell comparison’* we used Fisher’s Exact Test to look for differences between groups in the ratio of ants that circumvented or fell into the trap. For the ‘*Scan number and meander’* comparisons ants were followed individually across successive trials. We compared *scan number* and *meander* across three situations: (i) Trial before trap; (ii) Trial with first trap-fall; (iii) Trial following first trap-fall for two sections of the route, before the trap and after the trap, separately. We used a generalised linear mixed effects model (glme) with ants as a random effect, followed by a Tukey’s test. We assumed a Poisson distribution for scan number (count data per ant) and a normal distribution for meander values.

## Acknowledgment

We would like to thanks Cody Freas and Jeanne Delor for help on the field, as well as the Centre for Appropriate Technology Limited (CfAT Ltd, Alice Springs) for field facilities and letting us running the experiment on their grounds.

---

**Figure S1.**
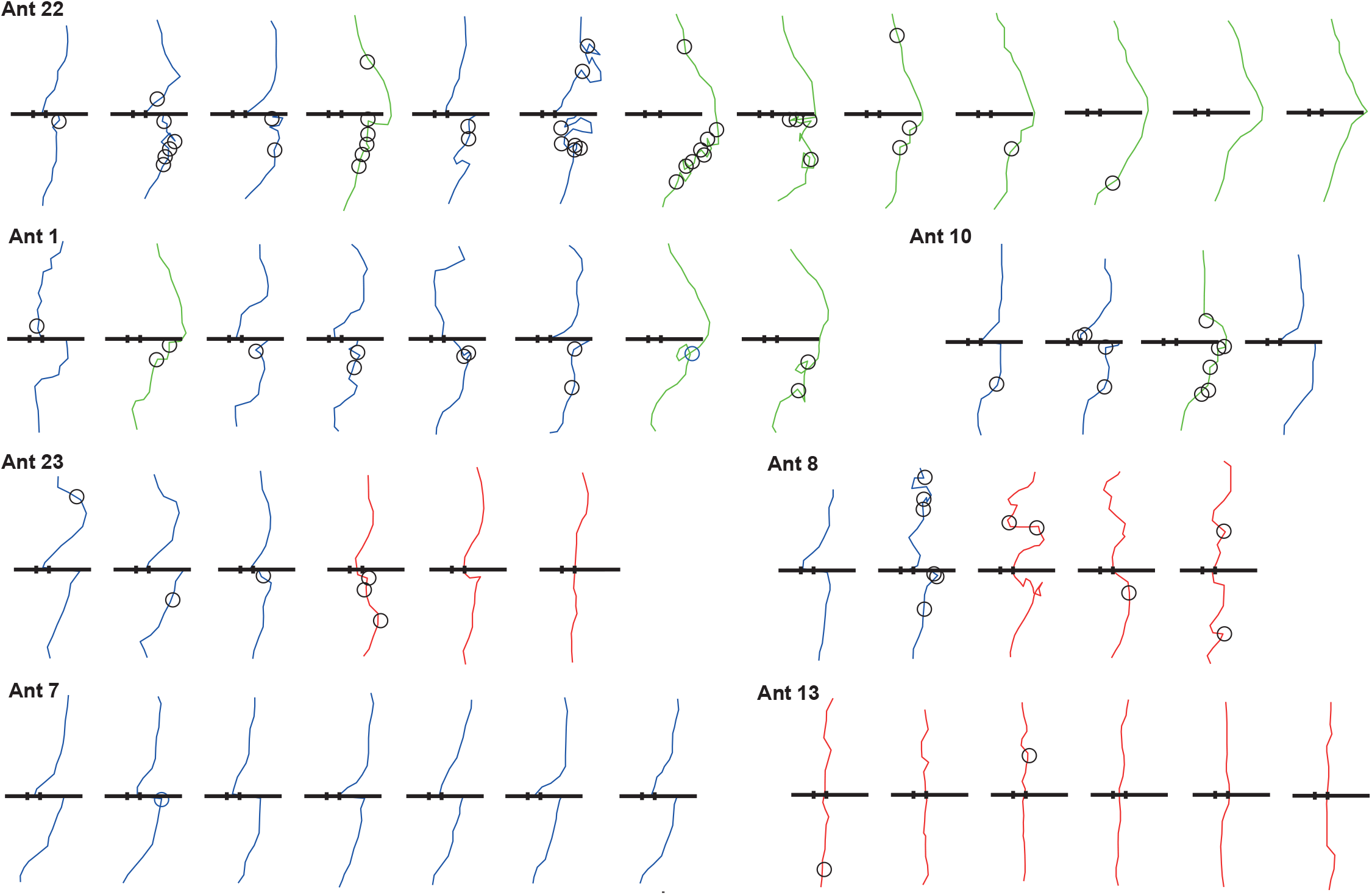
Route shape and scanning ontogeny for individual ants. For individual ants we show successive routes from the first run incorporating the trap. Scan locations are marked with a circle and routes are colour coded as in Fig 1 and 2 with the addition of paths marked in red for ants that learnt to use the stick bridge efficiently.

